# P2X2 receptor subunit interfaces are missense variant hotspots where mutations tend to increase apparent ATP affinity

**DOI:** 10.1101/2021.03.26.436616

**Authors:** Federica Gasparri, Debayan Sarkar, Sarune Bielickaite, Mette Homann Poulsen, Alexander Sebastian Hauser, Stephan Alexander Pless

## Abstract

**Background and Purpose:** P2X receptors (P2XRs) are trimeric ligand-gated ion channels (LGICs) that open a cation-selective pore in response to ATP binding to their large extracellular domain (ECD). The seven known P2XR subtypes can assemble as homo- or heterotrimeric complexes and contribute to numerous physiological functions, including nociception, inflammation and hearing. The overall structure of P2XRs is well established, but little is known about the spectrum and prevalence of human genetic variations and the functional implications in specific domains.

**Experimental Approach:** Here we examine the impact of P2X2 receptor (P2X2R) inter-subunit interface missense variants identified in the human population or through structural predictions. We test both single and double mutants through electrophysiological and biochemical approaches.

**Key results:** We demonstrate that predicted ECD inter-subunit interfaces display a higher-than-expected density of missense variations and that the majority of mutations that disrupt putative inter-subunit interactions result in channels with higher apparent ATP affinity. Lastly, we show that double mutants at the subunit interface show significant energetic coupling, especially if located in close proximity.

**Conclusions and Implications:** We provide the first structural mapping of the mutational burden across the human population in a LGIC and show that the density of missense mutations is constrained between protein domains, indicating evolutionary selection at the domain level. Our data may indicate that, unlike other LGICs, P2X2Rs have evolved an intrinsically high threshold for activation, possibly to allow for additional modulation or as a cellular protection mechanism against overstimulation.

**Bullet point summary:**

‘What is already known’:

- P2X2 receptors are ATP-activated ion channels implicated in hearing and nociceptice pathways
‘What this study adds’:

- A structural mapping of missense variants observed in the human population
- We identify the intersubunit-interface as a variant hotspot and decipher functional impact of mutations
‘Clinical significance’:

- The development of both inhibitors and activators of P2X2 receptor function may be required

## INTRODUCTION

Release of ATP into the extracellular milieu can activate a class of trimeric ligand-gated ion channels known as P2X receptors (P2XRs) (Burnstock, 1972; Coddou et al., 2011). Upon ATP binding, P2XRs open a cation-selective pore and contribute to a variety of physiological processes. These include nociception, sensory transduction, inflammatory processes and muscle contraction, highlighting P2XRs as potential drug targets (Arulkumaran et al., 2011; Broom et al., 2008; Burnstock, 2007; Finger et al., 2005; Illes et al., 2020; Jarvis et al., 2002; Khakh & North, 2012). Seven P2XR isoforms (P2X1-7) are known in humans, and they display tissue-specific expression patterns and most subunits can assemble as homo- or heterotrimeric receptors. For example, homomeric P2X2 receptors (P2X2Rs) are involved in hearing (George et al., 2019; Zhu et al., 2017) and P2X2/P2X3 heteromeric receptors are implicated in nociceptive pathways (Carter et al., 2009; Honore et al., 2006; Stephan et al., 2018).

A number of recent structural studies have provided unprecedented insight into the three-dimensional architecture of P2XRs (Hattori & Gouaux, 2012; Karasawa & Kawate, 2016; Kasuya et al., 2016; Kasuya, Fujiwara, et al., 2017; Kasuya, Yamaura, et al., 2017; Kawate et al., 2009; Mansoor et al., 2016; McCarthy et al., 2019; Wang et al., 2018). Overall, the receptors adopt a chalice-shaped trimeric structure with each subunit roughly resembling the outline of a dolphin: the two helices (M1 and M2) of the transmembrane domain (TMD) form the fluke, while the large extracellular domain (ECD) makes up the body with attached dorsal fin, flippers and head domains (Kawate et al., 2009). The ATP binding site is located at the interface of two adjacent subunits and the contributions by conserved side chains or backbone atoms to the coordination of the ligand molecule have been investigated thoroughly (Chataigneau et al., 2013; Ennion et al., 2000; Gasparri et al., 2019; Jiang et al., 2000; Kasuya, Fujiwara, et al., 2017; Roberts et al., 2008; Roberts & Evans, 2006). Binding of ATP is thought to cause a series of conformational steps that ultimately trigger channel opening (Jiang et al., 2012; Roberts et al., 2012; Stelmashenko et al., 2014). Initially, a tightening of the jaw region around the ATP binding site causes a displacement of the surrounding flexible regions, dorsal fin and flippers (Jiang et al., 2011; Zhao et al., 2014). These movements exert tension on the β-sheet wall across upper and lower body, causing it to flex outward, enlarge the lateral fenestration present in the lower body region and, in turn, open the transmembrane pore (Chataigneau et al., 2013; Mansoor et al., 2016).

Although the consequences of mutations related to ATP binding and agonist-induced conformational changes are well-documented, it remains unclear where and to what extent human genetic variations are present in the different P2X2R domains and if they result in functional consequences. This is relevant and timely because with the advent of large-scale exome and whole genome sequencing efforts, numerous amino acid-altering missense variations have been identified and individual mutations in various protein families have been implicated in disease states (Stefl et al., 2013), cancer progression (Kamburov et al., 2015) or altered drug response (Hauser et al., 2018).

Here, we performed an *in silico* analysis for genetic variant hotspots and establish that among the different (sub)domains of the P2X2R, the inter-subunit interface shows the highest frequency of missense mutations, while the ATP binding site and the TMDs are least affected. Motivated by this finding, we focused primarily on mutational disruptions of interactions formed at the extracellular subunit interfaces in rat P2X2R. We show that interfering with putative inter-subunit interactions results in an increase in apparent ATP affinity, as about 80 % of the examined mutants displayed a significantly *reduced* EC_50_ for ATP compared to wild-type (WT). We further use double-mutant cycle analysis to demonstrate that the majority of tested sites show strong energetic coupling, thus revealing a tight interplay between residues throughout the ECD. Together, our data demonstrate that inter-subunit interactions are crucial for fine-tuning ATP sensitivity and, unusually, may contribute towards lowering the apparent agonist affinity of P2X2Rs.

## METHODS

### Modelling, genetic variation data, and conservation analysis

A human P2X2 homology model was built using SWISS-MODEL based on the human ATP-bound open state P2X3 structure [PDBid: 5SVK] (Bienert et al., 2017; Mansoor et al., 2016). Sequence identity was determined at 50.85% with an average model confidence of 0.68 ± 0.05. Types of interactions in the interface between subunits have been determined by PDBePISA (https://www.ebi.ac.uk/pdbe/prot_int/pistart.html) collectively rendering the “interface” (Schlee et al, 2019). Other domains such as the intracellular domain (ICD), transmembrane domain (TMD), and extracellular domain (ECD) have been determined from DSSP secondary structure prediction in PyMol (see Table S1). “ATP” has been defined as residue positions that are within 5 Å of ATP, annotated from *Chataigneau et al.* as resulting in ‘major decrease on ATP potency’ (i.e. at least 5-fold decrease) or ‘non-functional’ receptors (Chataigneau et al., 2013).

We considered the gene for the hP2X2R to be located on chromosome 12:132,618,776-132,622,388 on the forward strand spanning 11 exons with all variant alleles in reference to the Ensembl canonical transcript ENST00000643471.2. We obtained natural genetic variation data for P2X2 (ENST00000643471) from the Genome Aggregation Database (gnomAD) v. 3.1.1 (obtained on 11/05/2021), which compiled data on 125,748 exome- and 15,708 whole-genome sequences from aggregated human sequencing studies spanning seven human populations (Karczewski et al., 2020). We excluded singleton variants from our analysis to avoid potential biases from sequencing errors. These are expected to be specifically enriched in the error rate and can dramatically impact the false positive rate for rare variants, especially for larger sample sizes (Johnston et al., 2015; Ma et al., 2019). All the non-singleton missense variants have been resampled across the protein sequence, i.e. protein sequence positions were randomly assigned for the number of observed variants (82 unique variant positions for the ∼140,000 individuals) to arrive at the expected sampling distribution under random conditions. We performed 100,000 permutations and calculated the z-score and p-values (two-tailed) relative to the sampled mean and standard deviation for each domain.

Conservation scores have been calculated by ConSurf using Bayesian method based on a MAFFT alignment of 150 sequences from UNIREF90 using default parameters (Ashkenazy et al., 2016; Katoh & Standley, 2013). Predicted deleteriousness for each variant were determined and obtained by combined annotation-dependent depletion (CADD) (https://cadd.gs.washington.edu) (Rentzsch et al., 2019). Mean conservation scores and CADD deleteriousness scores were aggregated for each P2X2 domain.

### Chemicals

Adenosine 5′-triphosphate, disodium salt, hydrate (ATP, purity 99%) and salts were from Sigma-Aldrich.

### Mutagenesis and expression of P2X2R in *Xenopus laevis* oocytes

Point mutations were introduced in the cDNA of the rat P2X2 receptor (rP2X2R (P49653-1), sub-cloned into the pNKS2 vector) via PCR with custom-designed primers (Eurofins Genomics, Sigma-Aldrich) and *Pfu*Ultra II Fusion HS DNA polymerase (Agilent Technologies). Generally, positively charged amino acids (R and K) were substituted with Q; acidic residues (E and D) were mutated to Q, N or A; A was introduced instead of V, L or I; S was replaced by A and Y by F (to remove hydroxyl group) and H was mutated to L. The Ambion mMessage mMACHINE SP6 transcription kit (Thermo Fisher Scientific) was used to transcribe the rP2X2 cDNA to mRNA after linearization with Xho I (New England Biolabs). The reaction was purified with RNeasy columns (Qiagen) and mRNA was stored at – 80°C until use. Stage V-VI oocytes were surgically removed from *Xenopus laevis* frogs (anesthetized in 0.3% tricaine, according to license 2014-15-0201-00031, approved by the Danish Veterinary and Food Administration) and digested with collagenase (1.5 mg ml^−1^, Roche), dissolved in OR2 (82 mM NaCl, 2.5 mM KCl, 1 mM MgCl_2_, 5 mM HEPES adjusted to pH 7.4 with NaOH), under continuous shaking. Oocytes were incubated in OR2 at 18 °C and gently shaken until injection with mRNA. For electrophysiological recordings, WT and mutant rP2X2R mRNAs (concentration 50 - 3000 ng μl^−1^) were injected into the oocyte cytoplasm with a Nanoliter 2010 injector (World Precision Instruments). The volume of mRNA injected varied depending on the construct (10-50 nl). Injected oocytes were incubated in OR3 solution (Leibovitz’s L-15 medium (Life Technologies) with 5 mM L-glutamine, 2.5 mg ml^−1^ gentamycin, 15 mM HEPES, pH 7.6 with NaOH) or ND96 solution (in mM: 96 NaCl, 2 KCl, 1.8 CaCl_2_, 1 MgCl_2_, 5 HEPES, pH 7.4) supplemented with 2.5 mM sodium pyruvate, 0.5 mM theophylline, 0.05 mg ml^−1^ gentamycin and 0.05 mg ml^−1^ tetracycline and gently shaken at 18 °C until the day of the experiment.

### Electrophysiological recordings and data analysis

One to two days after mRNA injection, oocytes were transferred into a recording chamber (Dahan et al., 2004) and continuously perfused with Ca^2+^-free ND96 solution (in mM: 96 NaCl, 2 KCl, 1.8 BaCl_2_, 1 MgCl_2_, 5 HEPES, pH 7.4) through an automated, gravity-driven perfusion system operated by a ValveBank™ module (AutoMate Scientific). ATP solutions were freshly made prior to recordings. Solutions were prepared from agonist stocks (10 or 100 mM, stored at - 20°C) or directly weighted out and dissolved in ND96 to the desired final concentration (pH adjusted to 7.4 with NaOH). Oocytes were clamped at −40 mV and 10 sec applications of increasing concentration of ATP were used to activate rP2X2Rs. Between each application, oocytes were perfused for 1 min with ND96 to allow for full recovery from desensitization. Currents were recorded using borosilicate microelectrodes (Harvard Apparatus, resistance of 0.3-2 MΩ), backfilled with 3 M KCl, and an Oocyte Clamp C725C amplifier (Warner Instrument Corp.). Current was acquired at 500 Hz and digitized via an Axon™ Digidata® 1550 interface and the pClamp (10.5.1.0) software (Molecular Devices). The signal was further digitally filtered at 10 Hz with an eight-pole Bessel filter for analysis and display. ATP-elicited concentration-response curves (CRCs) were obtained by plotting the normalized peak current amplitude against the ATP concentration for each individual recording and subsequently fitted to the Hill equation in Prism v7 (GraphPad) to calculate EC_50_ values. These were averaged and reported as mean ± SD.

### Cell surface biotinylation and Western blots

Oocytes were injected with 9.2 nl of mRNA coding for rP2X2 WT (0.05 µg µl^−1^), 46 nl rP2X2-D78N-E167A (0.9 µg µl^−1^), 46 nl rP2X2-E91Q-R313Q (0.9 µg µl^−1^) and 9.2 nl rP2X2-Y86F-L276A (0.05 µg µl^−1^). Following incubation in OR3 solution for 36 hrs at 18°C, the oocytes were washed twice with PBS-CM (in mM: 137 NaCl, 2.7 KCl, 10 Na_2_HPO_4_, 1.8 KH_2_PO_4_, 0.1 CaCl_2_, 1 MgCl_2_) and surface proteins from 50 oocytes per construct were labeled using EZ-Link™ Sulfo-NHS-SS-Biotin (Thermo Fischer Scientific), dissolved to a final concentration of 1.25 mg ml^−1^ in ice cold PBS-CM. Following agitation for 30 min, the reaction was quenched for 30 min using quenching buffer (PBS-CM supplemented with 200 mM glycine). Next, the oocytes were lysed using lysis buffer (150 mM NaCl, 100 mM Tris-HCl, 0.1% SDS and 1% Triton-X-100) with added Halt protease inhibitor cocktail (1:100) (Thermo Fisher Scientific). Biotin-labeled surface proteins were isolated and purified using Pierce™ Spin Columns - Snap Cap (Thermo Fischer Scientific) with 500 µl of Pierce™ NeutrAvidin™ Agarose added to each column (Thermo Fisher Scientific). Purified surface proteins or total cell lysates were separated on a NuPage 3-8 % Tris-acetate protein gel (Thermo Fisher Scientific) at 200 V for 40 min and transferred to a PVDF membrane. Membranes were incubated in LI-COR blocking buffer for 1 hr, followed by incubation in LI-COR blocking buffer containing rabbit polyclonal anti-P2X2 (#APR-003, Alomone labs; 1:2000) and mouse anti-Na^+^/K^+^-ATPase (05-369; EMD Merck Millipore) at 4 °C overnight. Dilutions of secondary antibody was prepared for each blot. Next, the membranes were washed 5×2 min with TBST (20 mM Tris-HCl, 150 mM NaCl, 0.1% Tween 20, pH 7.5) and incubated with secondary antibodies (IRDye 800CW goat-anti-rabbit (1:5000, 925-32211; LI-COR Biosciences) and IRDye 680RD goat-anti-mouse (1:5000, 926-68070; LI-COR Biosciences) in LI-COR blocking for 1 hr at RT in the dark. Finally, the membranes were washed 5×2 min in TBST before being imaged using a PXi gel imaging station (Syngene). The western blot protocol described above and the data provided adhere to guidelines advised in *Alexander et al.* (Alexander et al., 2018).

### Double-mutant cycle analysis

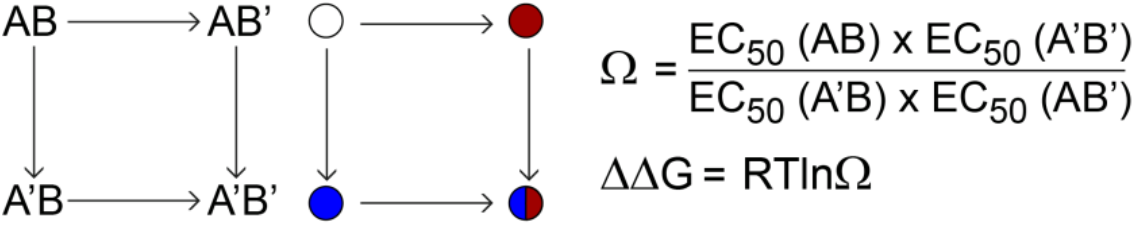

Scheme illustrating the principle of double mutant cycle analysis; AB represents WT protein, AB’ and A’B- two different single mutants, A’B’- protein containing both mutations. Coupling coefficient Ω for the two residues- A and B- is calculated from the apparent EC_50_ values of WT, double mutant, and single mutants. Coupling energy (or free energy change) ΔΔG is further calculated from coupling coefficient Ω, the gas constant R (8.314 J mol^−1^. K^−1^), room temperature T (298 K) (Horovitz, 1996; Schreiber & Fersht, 1995). **Statistical Analysis**

Statistical analysis was performed using Prism v7, significant differences were determined by performing unpaired t-test with Welch’s correction to a control value (i.e WT). For figure display, a single fit to the average normalized response (± SD) is shown using Prism v7 (Graphpad). Each P2X2R variant evaluated for concentration response analysis was tested in at least 5 oocytes from a minimum of two batches of cells. A consistent probability margin was used to define the threshold for level of significance while comparing mutants (**, *p* < 0.01; ***, *p* < 0.001; ****, *p* < 0.0001). The experimental design, execution and data analysis comply with the guidelines of the British Journal of Pharmacology (Curtis et al., 2018).

## RESULTS

### Human population data reveals increased missense mutation density at inter-subunit interfaces

To assess the mutational burden observed in hP2X2Rs across an ostensibly healthy and unrelated human population, we obtained natural genetic variation data on amino acid-changing missense mutations from the Genome Aggregation Database (gnomAD) of around 140,000 exome and whole-genomes sequences across seven sub-populations (Karczewski et al., 2020). In total, we identified 203 unique missense variants across 163 residue positions of the hP2X2R with a mean minor allele frequency of 4.03 x 10^−5^. We excluded all singleton variants (variants seen only once in the data set) and thereby focused our analysis on the 91 variants across 82 positions with at least two allele counts to account for biases from potential sequencing errors (Johnston et al., 2015; Ma et al., 2019). Since there is no hP2X2R structure available, we mapped all 82 variant positions on a homology model based on the human ATP-bound open state hP2X3R structure [PDBid: 5SVK] (Mansoor et al., 2016) (Figure 1A). In order to test if any subdomains or regions of P2X2Rs are subject to a particularly high mutational burden, we classified P2X2R residue positions into the extracellular domain (ECD), transmembrane domain (TMD), intracellular domain (ICD), ATP binding site (ATP) and inter-subunit interface positions (interface) (Figure 1B). To this end, we identified 12 positions that have previously shown significant decrease in ATP potency (Chataigneau et al., 2013) as the ATP binding site (ATP). We then defined interface positions as residues in the ECD that were involved in at least one hydrogen bond or salt bridge between subunits. This resulted in 23 residues collectively referred to as the interface (Figure 1A; Table S1).

**Figure 1:**
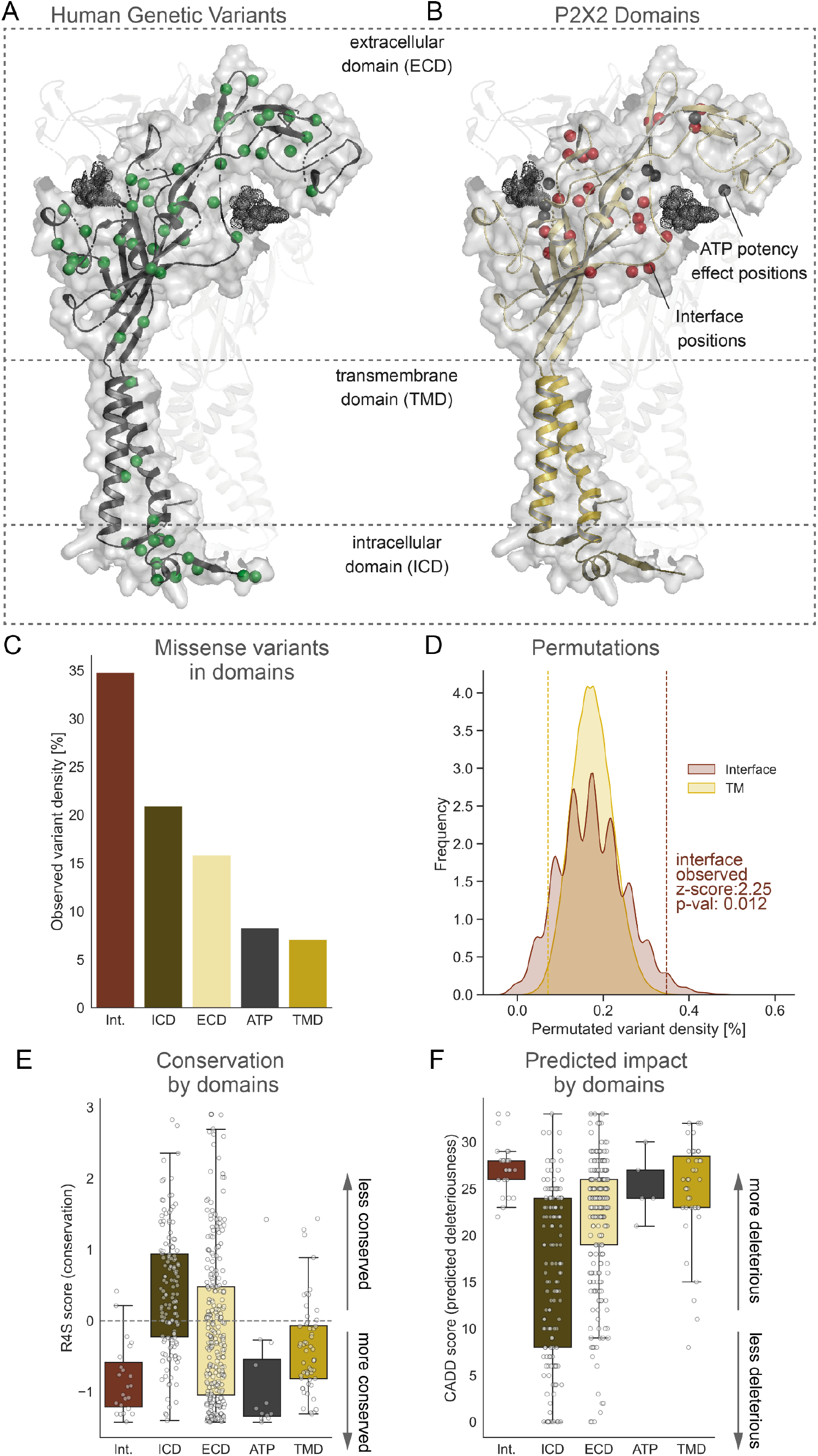
Human genetic variations of the P2X2 receptor. (A) P2X2 domains including ECD, TMD, ICD, Interface positions, and ATP potency effect positions annotated from literature (Chataigneau et al., 2013). (B) Distribution of all human genetic variations found among ∼140,000 individuals. Depicted structure displays a human P2X2 homology model based on the human ATP-bound open state P2X3 structure [PDBid: 5SVK] (Mansoor et al., 2016). (C) Density of missense variants (relative distribution by domain length). Note that ECD includes both ATP and interface positions. (D) Distribution of variant densities across 100,000 sampling simulations highlighting that the interface domain displays more genetic variants in the human population than expected from a random distribution of missense variants. (E) Evolutionary ortholog conservation by domain. (F) Predicted deleteriousness (CADD scores) by domain.

To evaluate a potential increase or decrease in mutational burden in any of these domains, we calculated the variation density, i.e. the fraction of positions that display variant carriers for each previously defined domain and functional region including ICD, TMD, ECD, ATP and the interface. For instance, among the 23 interface positions, we identified eight positions with missense variations (excluding singletons) among all individuals included (Table S2). Among the tested domains, the interface displays the highest variant density (∼35%), followed by intermediate values for the ICD (∼21%) and ECD (∼16%), while low variation was observed in the ATP binding site positions (∼8%) and the TMD (∼7%) (Figure 1C).

To further corroborate the high variation density in the interface domain, we employed a permutation test with 100,000 “mutation outcome simulations” to estimate the deviation from the mean of random expectation. From the random distribution, we computed the Z-score, which captures the distance of the actual number of observations (i.e. mutations in the interface domain) to the mean of random expectation in terms of the number of standard deviations. We estimated p-values as the ratio of the number of simulations where the random observations were greater than or equal to the number of actually observed values to the total number of randomizations. For the interface, our analysis predicts a mean variant density of 17% compared to the actually observed 35% (z-score: 2.252; p-val: 0.012), which translates into a significantly higher number of variants than expected from a random distribution (Figure 1D). By contrast, we predict a similar sampled variant density (17.4%) for the TMD, but in fact observe much lower densities (7%, z-score: 2.162; p-val: 0.015), resulting in far fewer variants than expected (Figure 1D). This could potentially indicate negative selection for mutations in the TMD region. At the same time, mean conservation of residues at the interface is slightly higher than for residues in the TMD (Figure 1E), and the predicted deleteriousness is highest across all domains and functional regions (Figure 1F).

In summary, we observe a much higher density of variants at the interface across our sample population than what would be expected from a random distribution. Next, we therefore sought to assess the functional impact of mutational disruptions at the inter-subunit interface.

### Functional impact of mutational interface disruptions based on human population data

To disrupt side chain-mediated interactions and also account for possible non-identical side chains at equivalent positions in different P2XR isoforms and orthologs, we decided to replace small hydrophobic and hydrophilic amino acids (valine, leucine, isoleucine, serine, asparagine, aspartate) with alanine, larger charged amino acids such as arginine, lysine and glutamate with glutamine and tyrosine with phenylalanine throughout the majority of the study (notable exceptions are G92R and R28C, see below). The resulting rP2X2R mutants were expressed in *Xenopus laevis* oocytes and currents in response to ATP application were measured using two-electrode voltage-clamp (TEVC). The analysis of the resulting concentration-response curves (CRCs) revealed that E63A, R274Q, S284A and R313Q (rP2X2R numbering) all showed *increased* apparent ATP affinity (Figure 2 and Table 1). Only Y294F and G92R were less sensitive to activation by ATP than WT, although the interpretation of the latter with regards to the interface contribution is somewhat ambiguous because we are unable to predict if introduction of the large and positively charged arginine results in a side chain pointing towards or away from the interface. Further, and consistent with previous work, the CRC analysis of mutations at interface side chains directly involved in ATP binding and subunit assembly (N288 and R304, (Chataigneau et al., 2013)) showed drastically reduced apparent ATP affinity (Figure S1).

**Figure 2:**
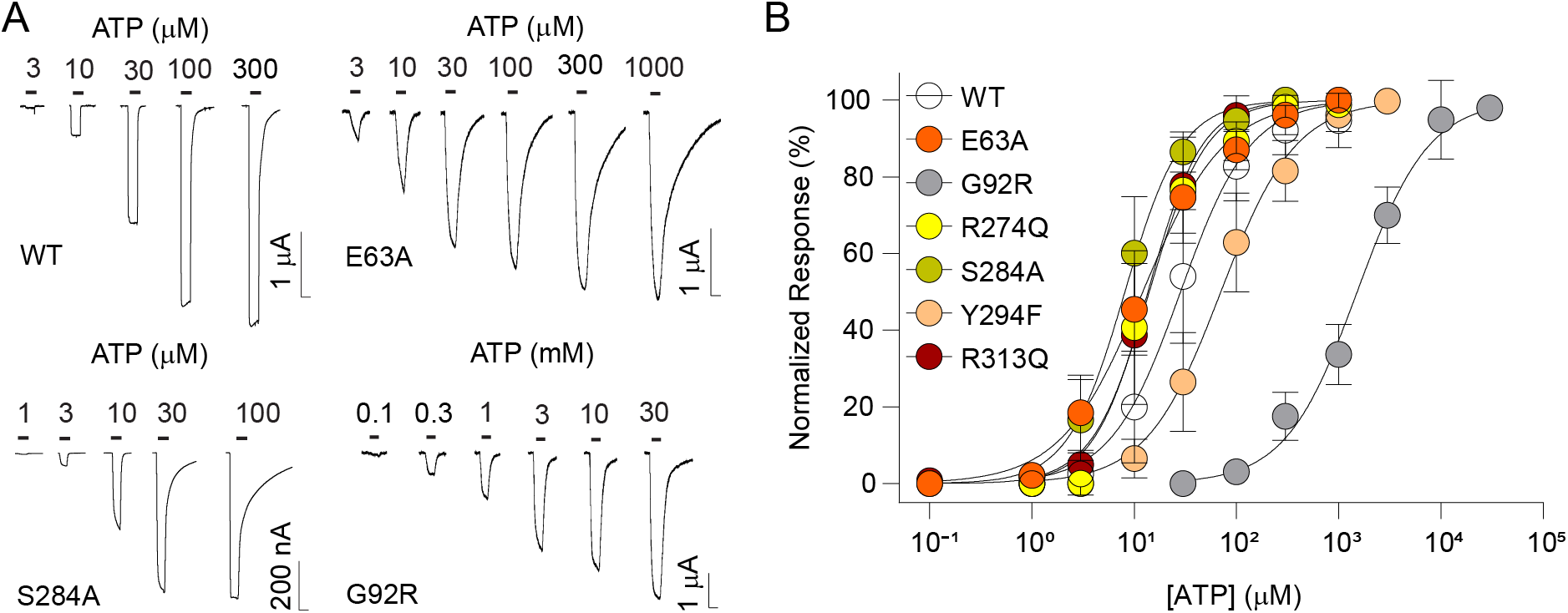
Functional impact of mutational interface disruptions based on human population data. (A) Example recordings of rP2X2 WT, E63A, S284A and G92R receptors. Currents are elicited by application of increasing concentrations of ATP (black bars). Scale bar: x, 10 s y, nA/µA. (B) Normalized ATP-elicited concentration-response data for WT and indicated mutant rP2X2Rs in response to application of increasing concentrations of ATP. Data are displayed as mean ± S.D. (n = 6-33).

**Table 1:** ATP-elicited concentration-response data (EC_50_) shown as mean ± S.D. as well as number of experiments (n) for WT and single mutants in the rP2X2R interface based on human population data (along with corresponding residue positions in hP2X2 and zfP2X4 receptors). Significant differences were determined by unpaired t-test. **, *p* < 0.01; ***, *p* < 0.001; ****, *p* < 0.0001.

| rP2X2 construct | Corresponding hP2X2 residue | Corresponding zfP2X4 residue | $EC_{50} \pm SD$ ($\mu M$) | n |
| --- | --- | --- | --- | --- |
| WT | N.A. | N.A. | $31.0 \pm 14.5$ | 33 |
| R28C | R40 | K30 | $29.3 \pm 17.5$ | 10 |
| E63A | E75 | L64 | $12.8 \pm 4.2$**** | 8 |
| G92R | G104 | E98 | $1590.9 \pm 443$**** | 9 |
| R274Q | R286 | R280 | $15.4 \pm 5.8$*** | 6 |
| S284A | S296 | A292 | $9.1 \pm 4$**** | 10 |
| N288A | N300 | N296 | $2566.6 \pm 502$**** | 7 |
| Y294F | Y306 | Y302 | $76.9 \pm 32.5$ ** | 8 |
| R304Q | R315 | R312 | $3011.8 \pm 862.4$**** | 7 |
| R313Q | R324 | R321 | $13.7 \pm 3$**** | 9 |

### Functional impact of mutational interface disruptions based on computational predictions

An obvious caveat of our above interface analysis is the fact that it was based on the hP2X3 structure because we do not have access to the structure of a P2X2 receptor. This creates a degree of ambiguity with regards to the positions identified as interface positions. We therefore set out to identify inter-subunit interface positions based on a distinct and independent approach. To this end, the apo and ATP-bound zebrafish P2X4R structures (PDBid 315D and 4DW1 (Hattori & Gouaux, 2012; Kawate et al., 2009)) were analyzed using the PISA (Protein, Interfaces, Structures and Assemblies (EMBL-EBI, n.d.)) program to pinpoint residues at the interface between P2XR subunits that form H-bonds and/or salt bridges in both conformational states (Table S3, see also (Hausmann et al., 2014)). To evaluate if these interactions are conserved in P2X2Rs, we used a sequence alignment to identify the corresponding residues in the rP2X2R subtype. Similar to the above, we excluded side chains at the ATP binding pocket, which have been extensively characterized previously (Chataigneau et al., 2013; Gasparri et al., 2019; Jiang et al., 2000), and instead focused on mutating residues that would disrupt putative inter-subunit interactions distant from the ligand binding pocket. This led to the identification of 18 sites, only two of which (S284 and R313) overlapped with the hP2X3-based analysis outlined above. Analysis of CRCs showed that 12 of the 15 newly designed single mutants (Figure 3) responded to lower concentrations of ATP than WT rP2X2R, resulting in significantly reduced EC_50_ values (Figure 4A-D, Table 2). Six of these 12 mutants (S65A, E84Q, S190A, L276A, K293Q, Y295F) even showed *∼*10-fold decrease in EC_50_, while K79Q did not change the EC_50_ compared to WT and I73A and V80A both resulted in about two- and ten-fold increase in EC_50_ (Table 2).

**Figure 3:**
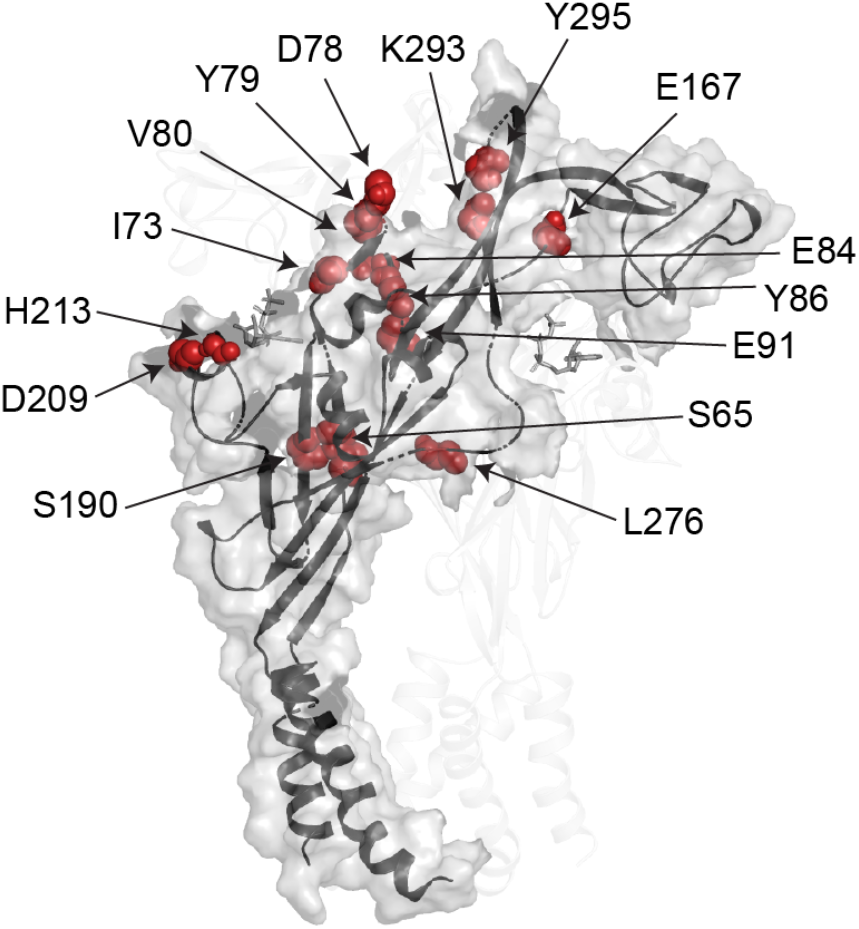
Homology model of rP2X2R based on the ATP-bound zebrafish P2X4R structure (PDBid 4DW1 (Hattori & Gouaux, 2012) with interface residues identified using PISA shown as red spheres.

**Figure 4:**
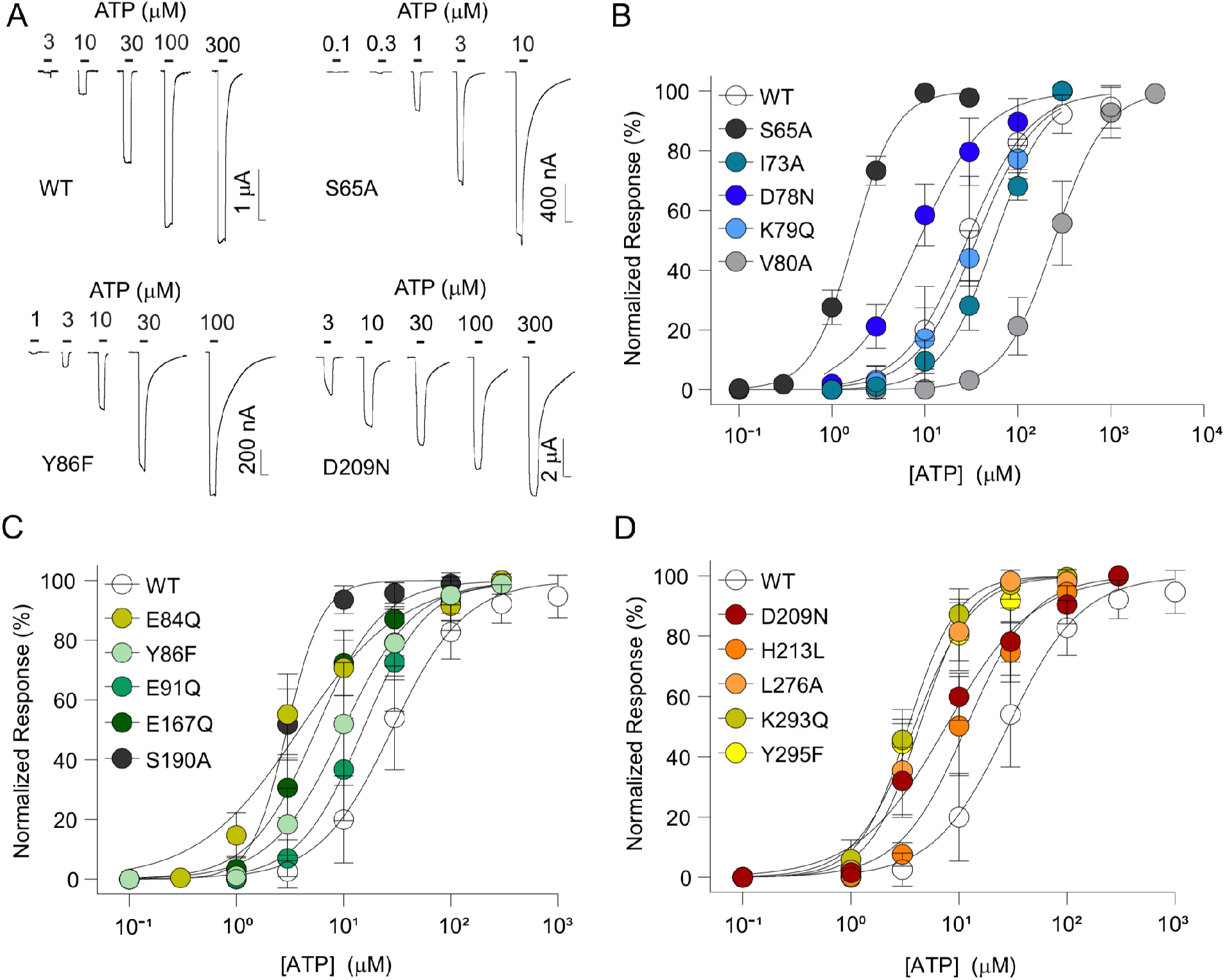
Characterization of single point mutations at the inter-subunit interfaces of rP2X2R. (A) Example recordings of rP2X2 WT, S65A, Y86F and D209N receptors. Currents are elicited by application of increasing concentrations of ATP (black bars). Scale bar: x, 10 s y, nA/µA. (B)-(D) Normalized ATP-elicited concentration-response data for WT and indicated mutant rP2X2Rs in response to application of increasing concentrations of ATP. Data are displayed as mean ± S.D. (n = 6-33).

**Table 2:** ATP-elicited concentration-response data (EC_50_) shown as mean ± S.D. as well as number of experiments (n) for WT and single mutants lining the P2X2R inter-subunit interface (along with corresponding residue positions in hP2X2 and zfP2X4 receptors). Significant differences were determined by unpaired t-test. **, *p* < 0.01; ***, *p* < 0.001; ****, *p* < 0.0001; ^a^, residues in the TM domain of P2X2R; ^b^, residues away from the interface (S122, T123) or previously characterized (E167 in (Hausmann et al., 2013)).

| rP2X2 construct | Corresponding hP2X2 residue | Corresponding zfpP2X4 residue | $EC_{50} \pm SD$ (μM) | n |
| --- | --- | --- | --- | --- |
| WT | N.A | N.A. | $31.0 \pm 14.5$ | 33 |
| Y43A<sup>a</sup> | Y55 | Y45 | $0.7 \pm 0.3$**** | 6 |
| Y43F<sup>a</sup> | Y55 | Y45 | $11 \pm 2$**** | 5 |
| S65A | S77 | S66 | $1.7 \pm 0.2$**** | 6 |
| I73A | I85 | I74 | $56 \pm 12$** | 6 |
| D78N | H90 | E84 | $9.3 \pm 3$**** | 11 |
| K79Q | K91 | R85 | $36 \pm 10$ | 6 |
| V80A | V92 | I86 | $263.2 \pm 100.5$*** | 8 |
| E84Q | E96 | A90 | $4.1 \pm 2$**** | 9 |
| Y86F | Y98 | Y92 | $11 \pm 5$**** | 10 |
| E91Q | E103 | Q97 | $17 \pm 7$**** | 12 |
| S122C<sup>b</sup> | A134 | S127 | $3.4 \pm 1$**** | 6 |
| T123C<sup>b</sup> | T135 | T128 | $1.9 \pm 0.3$**** | 6 |
| E167Q<sup>b</sup> | E179 | E171 | $6.1 \pm 2$**** | 8 |
| S190A | S202 | N195 | $3 \pm 0.4$**** | 7 |
| D209N | G221 | S215 | $8.2 \pm 3$**** | 6 |
| H213L | R225 | H219 | $13 \pm 5$**** | 6 |
| L276A | L288 | L282 | $4.5 \pm 1$**** | 12 |
| K293Q | K305 | K301 | $3.5 \pm 0.9^{****}$ | 10 |
| Y295F | Y307 | Y303 | $4.1 \pm 1^{****}$ | 6 |

Next, we set out to assess if inter-subunit residues in the transmembrane domain (TMD) would also affect the ATP EC_50_ value, as for example suggested by the constitutively active phenotype of a P2X2R M1 mutation involved in hearing loss (V60L in hP2X2R) (George et al., 2019). The hydroxyl moiety of the tyrosine in position 43 of rP2X2R points towards the M2 helix of the adjacent subunit and single point mutations to either alanine or phenylalanine resulted in a pronounced left-shift in the ATP CRCs compared to the WT (Table 2).

This indicates that disruptions to inter-subunit interactions in both the TMD and in the ECD, not involved in ATP binding, are crucial for channel gating. Interestingly, mutating single residues most often resulted in phenotypes with increased apparent ATP affinity.

### Most double mutants at the same subunit interface show energetic coupling

Next, we sought to investigate if introduction of double mutants at the inter-subunit interface would result in additive effect in rP2X2R apparent ATP affinity. In a trimeric channel, there are three equivalent subunit interfaces within each fully assembled receptor. Here, we first generated a series of double mutants in which each of the inter-subunit interfaces was disrupted by two of the above characterized single mutations situated on an interface facing the *same* subunit. Specifically, we generated the S284A/L276A, Y86F/K293Q, E84Q/E91Q, D78N/S190A and V80A/S190A mutations (Figure 5A).

**Figure 5:**
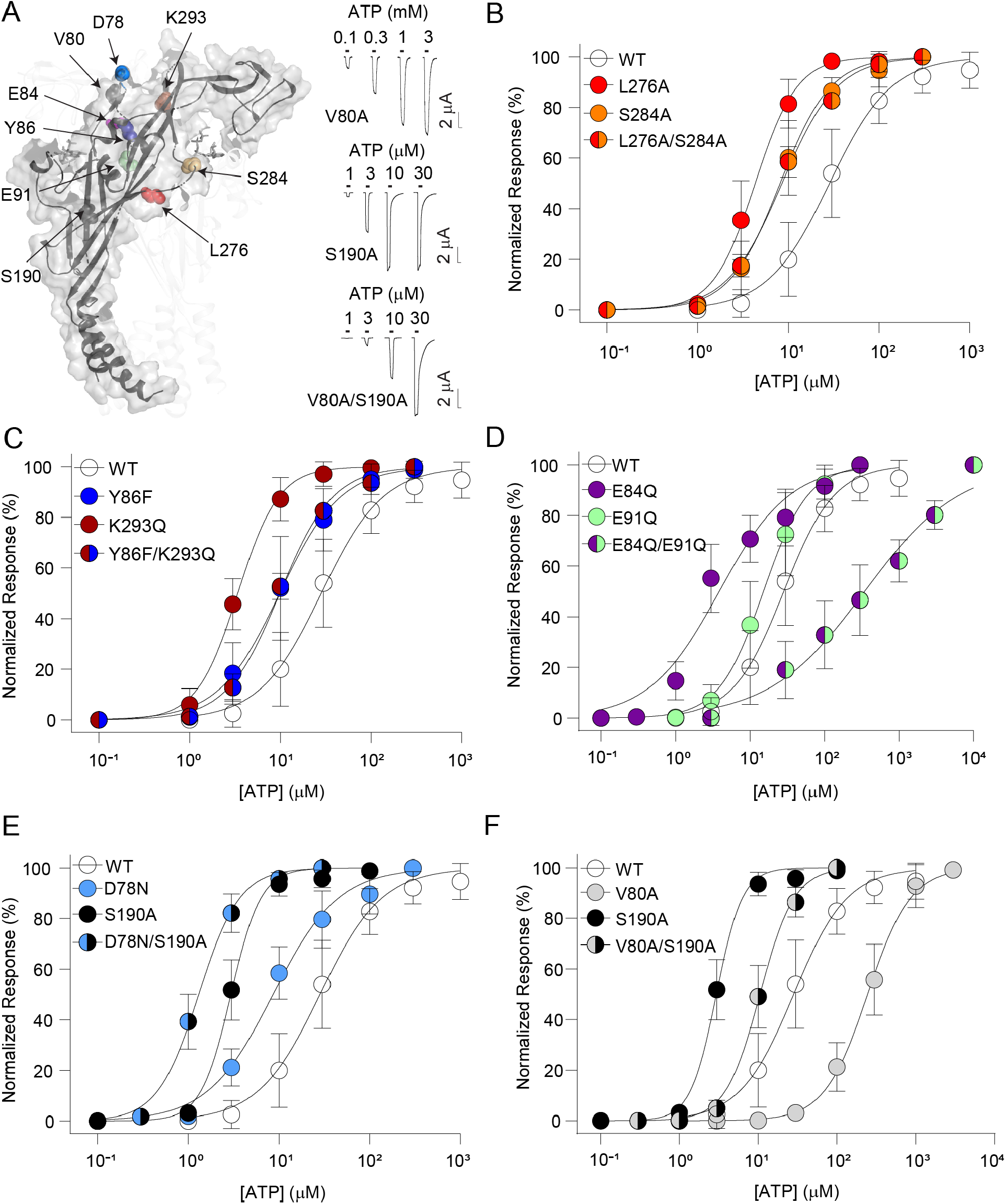
Characterization of double mutants disrupting the same subunit interface. (A, left) Homology model of rP2X2R with residues corresponding to the positions mutated shown as spheres. (A, right) Example recordings of V80A, S190A and V80A/S190A mutants. Currents are elicited by application of increasing concentrations of ATP (black bars). Scale bar: x, 10 s y, µA. (B) to (F) Normalized ATP-elicited concentration-response data for WT (empty symbols), single (single-colour symbols) and double mutants (split-colour symbols) rP2X2Rs in response to application of increasing concentrations of ATP. Data are shown as mean ± S.D. (n = 5-33).

Both L276 and S284 are situated on the left flipper region and, when mutated to alanine, reduced the EC_50_ values to < 10 µM (Figure 5B and Table 3). If the two mutations were energetically uncoupled, we would expect the effects on the EC_50_ to be additive, i.e. the S284A/L276A double mutant to display an even further reduced EC_50_ value. However, we determined the S284A/L276A double mutant EC_50_ to be indistinguishable from that of the S284A single mutant (Figure 5B). We calculated the coupling energy between the two mutants to be 4.6 kJ mol^−1^, indicating strong energetic coupling (Table 3). Despite being located distant from the S284A/L276A pair, the Y86F/K293Q double mutant in the upper body resulted in an almost identical coupling energy (5.2 kJ mol^−1^; Figure 5C, Table 3). Strikingly, the E84Q/E91Q double mutant pair (both positions are located between the β3-β4 sheets) displayed a 20-fold increase in EC_50_ compared to that of the WT, and in stark contrast to the reduced EC_50_s observed with the two single mutants (Figure 5D, Table 2 and 3). Our double-mutant cycle analysis revealed a very strong coupling of the E84/E91 pair (12.9 kJ mol^−1^).

**Table 3:** ATP-elicited concentration-response data (EC_50_) shown as mean ± S.D. as well as number of experiments (n) for WT and double mutants both at different and same subunit interface. Coupling energy values (kJ/mol) calculated as described in material and method section. Significant differences were determined by unpaired t-test. **, *p* < 0.01; ***, *p* < 0.001; ****, *p* < 0.0001; ^a^, residues located *away* from the subunit interface.

| rP2X2 construct | $EC_{50} \pm SD$ ($\mu M$) | n | Coupling energy, $\Delta\Delta G$, kJ/mol |
| --- | --- | --- | --- |
| WT | $31.0 \pm 14.5$ | 33 | ND |
| Y86F L276A | $12 \pm 9.8^{***}$ | 8 | 5.0 |
| D78N L276A | $18.6 \pm 7.9^{***}$ | 13 | 6.5 |
| D78N E167A | ND | ND | ND |
| E91Q R313Q | ND | ND | ND |
| L276A S284A | $8.6 \pm 1^{****}$ | 5 | 4.6 |
| Y86F K293Q | $10 \pm 1^{****}$ | 5 | 5.2 |
| E84Q E91Q | $414 \pm 269^{***}$ | 13 | 12.9 |
| S190A D78N | $1.3 \pm 0.3^{****}$ | 9 | 0.9 |
| S190A V80A | $11 \pm 3^{****}$ | 9 | -2.0 |
| L276A T123C<sup>a</sup> | $23.3 \pm 18.5$ | 19 | 11.0 |
| L276A S122C<sup>a</sup> | $34.4 \pm 23.4$ | 15 | 10.5 |
| Y86F T123C<sup>a</sup> | $21.6 \pm 9.8^{**}$ | 20 | 8.6 |

Next, we sought to interrogate potential upper-lower body interactions through different combinations of mutants at S190 (β8-sheet), D78 (β3-sheet) and V80 (β3-sheet). The D78N/S190A double mutant showed a lower EC_50_ compared to each of the underlying single mutants (Figure 5E, Table 3). Consequently, the change in free energy of approximately 0.9 kJ mol^−1^ indicated a small degree of coupling. Similarly, the V80A/S190A double mutant showed only modest change in free energy (−2.0 kJ mol^−1^; Figure 5F, Table 3).

Together, this suggests that double mutants within the flipper domain, as well as within the upper body show strong energetic coupling, while we did not observe such coupling between more distant upper-lower body pairs.

### Double mutants at different subunit interfaces show energetic coupling or prevent expression

Next, we generated a set of double mutants in which each of the inter-subunit interfaces was disrupted by two of the above characterized single mutations situated on an interface facing a *different* subunit. Specifically, three positions in the upper body D78, Y86 and E91 (in β3-β4 sheet and loop) were mutated in combination with side chains from the head domain (E167), left flipper (L276) or lower body (R313) to investigate if double mutations would be energetically coupled (Figure 6A).

**Figure 6:**
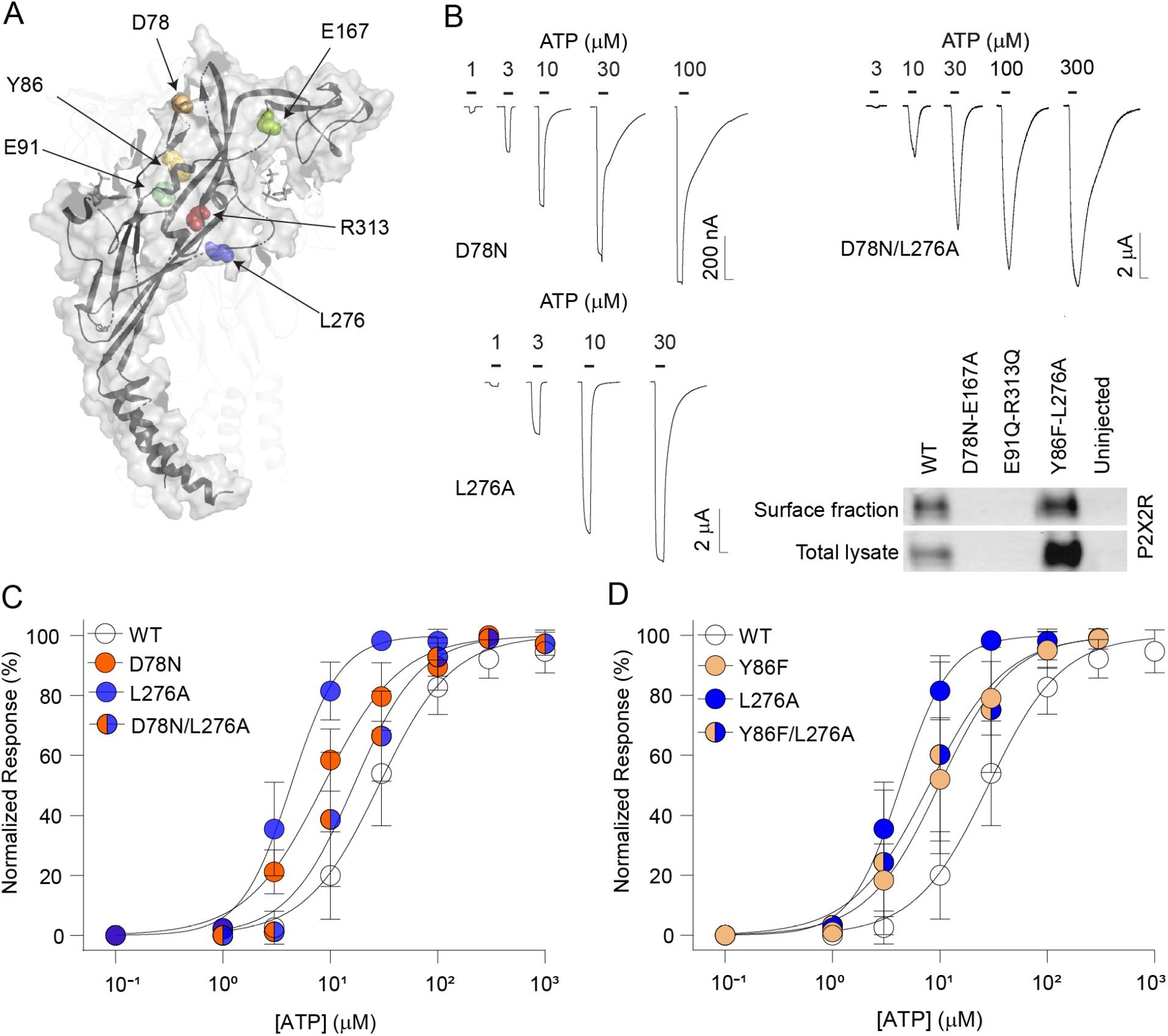
Characterization of double mutants disrupting different subunit interface. (A) Homology model of rP2X2R with residues corresponding to the positions mutated shown as spheres. (B) Example recordings of D78N, L276A and D78N/L276A and Western blot of surface fraction or total lysate extracted from oocytes expressing the indicated constructs (or uninjected oocytes). Note that the uncropped blot is provided as Figure S2. (C) and (D) Normalized ATP-elicited concentration-response data for WT (empty symbols), indicated single (single-colour symbols) and double mutants (split-colour symbols) P2X2Rs in response to application of increasing concentrations of ATP. Data are shown as mean ± S.D. (n = 5-33).

The E91Q/R313Q and D78N/E167A double mutants did not show any ATP-gated inward currents, even in response to high (10 mM) concentrations of ATP. In order to assess if this was due to severe gating phenotypes or rather surface expression, we performed a surface biotinylation assay, followed by Western blotting. As shown in Figure 6B, bands corresponding to a rP2X2R-sized protein are absent for the E91Q/R313Q and the D78N/E167A double mutant channels in both the surface fraction and the total lysate, suggesting that these double mutants are not expressed in *Xenopus laevis* oocytes.

By contrast, the Y86F/L276A and D78N/L276A double mutants displayed an EC_50_ similar to that observed with the single mutants, i.e. significantly left-shifted compared to WT rP2X2R (Figure 6C/D, Table 3). We performed double-mutant cycle analysis to assess a potential energetic coupling between these mutations. This yielded coupling energies of 5.0 kJ mol^−1^ for Y86F/L276A and of 6.5 kJ mol^−1^ for D78N/L276A (Table 3), suggesting strong energetic coupling between these side chain pairs.

### Energetic coupling with residues not lining the subunit interface

We then sought to assess if energetic coupling can also be observed for double mutants in which one of the mutations was located *away* from the subunit interface. We thus chose to generate the S122C and T123C single mutants in the head domain, which resulted in a pronounced left-shift in the ATP CRC (Table 2). Similarly, both the S122C/L276A and the T123C/L276A double mutants displayed a left-shifted EC_50_ compared to that of the WT and exhibited strong energetic coupling (10.5 kJ mol^−1^ and 11.0 kJ mol^−1^, respectively; Table 3). This was mirrored by the results obtained for the Y86F/T123C double mutant, which also showed high apparent ATP affinity (Table 3) and strong energetic coupling (8.6 kJ mol^−1^, Table 3). These findings suggest that pronounced energetic coupling is not unique to residues located at the subunit interface, but may be a more general property of the rP2X2R ECD.

### No measurable functional effects by a possibly clinically relevant P2X2 mutation

Lastly, we sought to identify P2X2 variants in aggregated PHEWAS data from the UK Biobank that could be associated with clinical traits (Canela-Xandri et al., 2018). Our analysis identified the genetic variant Arg40Cys in hP2X2 (rs75585377) at the bottom of TM1 to be associated with a number of blood phenotypes such as corpuscular volume (-log10(p-value): 15.23), reticulocyte volume (-log10(p-value): 13.08), corpuscular haemoglobin (-log10(p-value: 12.75) and sphered cell volume (-log10(p-value): 9.17). We found that the equivalent mutation in rP2X2 (R28C) had no significant effect on apparent ATP affinity (Fig S1 and Table 1). However, in light of the relatively lower degree of conservation in the N-terminus across P2X2R orthologs (both in terms of length and sequence identity), future studies on the hP2X2R and additional functional assays may be required to exclude possibly clinically relevant effects by the Arg40Cys variant in the hP2X2R.

## DISCUSSION

Analysis of the frequency of missense mutations in hP2X2Rs across an approximately representative sample of the human population reveals a strikingly uneven distribution of the mutational frequency. Both the ATP binding site and the TMD display very low mutational burden, likely due to their crucial role in P2X2R function and integrity. By contrast, our data reveal a surprisingly high number of missense mutations at the inter-subunit interfaces. This observation has potentially important implications because mutations at protein-protein interfaces often affect protein function and can be the cause of pathophysiologically relevant protein dysfunction (Iqbal et al., 2020; Jubb et al., 2017; Livesay & Marsh, 2021).

However, we did notice an apparent conundrum: on the one hand we identify more positions at the interface with missense variants in the human population than expected, while on the other hand, the interfaces display a high degree of ortholog conservation. There are multiple possible explanation for this observation: i) The variants we observe are variants without deleterious effects, potentially even with beneficial effects; ii) These variants are exceedingly rare and heterozygous; iii) Other positions at the interface, i.e. those without variants in the human population, are less susceptible for variations or iv) The positions we classify as interface from the various models are not overlapping exactly with the interface in the actual human protein.

Regardless of the origin for this phenomenon, the inter-subunit interface clearly stands out in our genetic analysis and we therefore embarked on a detailed functional investigation of mutations at sites predicted to be located at the interface. In absence of any P2X2 structure and to avoid bias, we pursued two distinct approaches to identify residues at the interface: first, we studied the impact of mutations at positions predicted to lie at the interface based on the hP2X3 structure (focusing only on sites that display two or more missense variants in the population) and, second, used an unbiased approach based on the zfP2X4 structure to identify interface-lining positions. Although only two sites overlapped between the two approaches (S284 and R313), numerous others were in very close proximity (E63 and S65; R274 and L276; Y294 and both K293 and Y295) and, strikingly, about 80% of mutations examined at the interface resulted in left-shifted CRCs (18 out of the total of 23). Together, this emphasizes the appropriateness of using two different and independent approaches and highlights an overall trend for functional outcomes of disruptions at the inter-subunit interface.

Mutations of conserved side chains within a protein of interest typically disrupt function. In the context of LGICs, this means that mutations in the ECD, including those at or near the subunit interface, tend to result in increased EC_50_ values for ligands binding at the orthosteric binding site or more generally disrupt channel function. This has been observed for a variety of LGICs, such as GlyR *α*1, nAChR *α*7 and iGluRs (Braun et al., 2016; Iacobucci et al., 2021; Tang et al., 2018; Tang & Lummis, 2018; Weston et al., 2006). In fact, even mutational scans in the ECD of a close cousin of the P2X2R, the P2X1R, have established that the vast majority of mutations result in increased EC_50_ values (Ennion et al., 2000; Roberts & Evans, 2004, 2006). Similarly, mutations in or near the ATP-binding pocket of a variety of P2XR subtypes have been shown to greatly increase EC_50_ values (Bodnar et al., 2011; Gasparri et al., 2019; Hausmann et al., 2013; Jiang et al., 2000; Roberts et al., 2008; Zemkova et al., 2007). Here, however, we find that about 80% of mutations designed to disrupt putative interactions across rP2X2R inter-subunit interfaces resulted in *lower* EC_50_ values. Importantly, this trend was independent of the chemical properties of the side chain in question, i.e. this was true for aromatic, hydrophobic and charged side chains. This finding is consistent with previous P2X2R studies, which demonstrated that individual mutations of side chains lining the subunit interface in the ECD or TMD result in lower EC_50_ values or even constitutive activity (George et al., 2019; Jiang et al., 2010; Jindrichova et al., 2009).

Given that many of the side chains mutated here are conserved across different P2XR isoforms (Kawate et al., 2009), it remains to be elucidated to what extent our findings apply to the other members of this receptor family. Also, in light of the lower EC_50_s observed for mutations away from the subunit interface (S122C and T123C in this study, E167R and H319A/K in work by others (Clyne et al., 2002; Hausmann et al., 2013; Sattler et al., 2020)), we cannot exclude the possibility that the trend observed for mutations at the subunit interface is not a more general feature of the rP2X2R ECD (outside the ATP binding pocket).

Finally, we sought to assess if combining two of the tested single mutants would indicate energetic coupling. To this end, we turned to double-mutant cycle analysis. Two of the mutant pairs (E91Q/R313Q and D78N/E167A) failed to express, but the remaining 10 pairs could be tested functionally (Table 3). Both double mutants involving S190A (S190A/D78N and S190A/V80A) showed no signs of strong coupling (ΔΔG values of 0.9 and −2.0 kJ/mol, respectively), possibly due to the large physical distance between the two mutations/residues. By contrast, we observed energetic coupling for the remaining double mutants we tested, especially for side chains in relatively close proximity within the structure (ΔΔG values >2.5 kJ/mol, see Table 3). Here, the E84Q/E91Q double mutant stood out in particular, with a ΔΔG value of 12.9 kJ/mol. This value is much higher than those observed in previous P2X2R studies (Hausmann et al., 2013; Jiang et al., 2010), but similar to that reported with side chains lining the ligand-binding site of a glutamate-gated chloride channel (Lynagh et al., 2017). Interestingly, strong energetic coupling was not restricted to double mutants along the subunit interface. In fact, the L276A-containing double mutants L276A/S122C and L276A/T123C both showed high ΔΔG values, although neither S122 and T123 are located at the subunit interface. This could suggest that residue pairs involving at least one interface side chain tend to be strongly coupled. However, it is also plausible that the strong coupling observed for the S122 and T123 mutants are due to more global disruptions caused by altered disulphide bond patterns in the cysteine-rich head domain (Lörinczi et al., 2012).

When interpreting our work, it is important to consider a number of limitations: i) we cannot exclude the possibility that the positions we classify as interface in this study may not overlap exactly with the interface in the actual human P2X2 receptor protein; ii) mutational effects could be masked or distorted because many inter-subunit interface side chains can engage in interactions with more than one other side chain or backbone; iii) our methodological approach is not capable of discerning all possible effects caused by the mutations. For example, it is unable to disentangle effects on binding and/or gating (Colquhoun, 1998). Further, the TEVC approach does not provide sufficiently high temporal resolution to address potential alterations in kinetics, changes in desensitization or changed membrane expression; iv) expression of rP2X2Rs in *Xenopus laevis* oocytes affords relatively high throughput, but functional and pharmacological properties may differ in mammalian cells or with the expression of the human clone. The latter two caveats are particularly relevant for the R28C mutation, which did not display a change in EC_50_ upon expression in oocytes, but may affect other receptor properties or show altered function in other cell types.

In conclusion, we find that mutations at the subunit interface of the rP2X2R ECD based on either hP2X2 receptor population data or homology model-derived data (based on the zfP2X4R channel structure), generally result in lower EC_50_ values. We further demonstrate that double mutations involving these sites typically show strong energetic coupling. This is true in particular for sites within close proximity, revealing a tight functional interplay between residues in the ECD. Although possibly not exclusive to inter-subunit locations or even ECD sites, these findings indicate that rP2X2Rs, unlike numerous other LGICs, have apparently not evolved for maximum agonist sensitivity. In support of this notion, their activation is fine-tuned by Mg^2+^, which when bound to ATP renders it a very ineffective agonist (Li et al., 2013). It is thus tempting to speculate that P2X2Rs have evolved towards low levels of activity, possibly as a cellular protection mechanism against overstimulation or as a means to enable additional modulation of agonist sensitivity. From a clinical perspective, this motivates the development of both P2X2R inhibitors and potentiators, in order to be able to eventually treat patient with mutations that either increase or decrease apparent ATP affinity.

## Abbreviations

ATP: adenosine 5′-triphosphate
PDB: protein data bank
CADD: Combined Annotation-Dependent Depletion
CRC: concentration-response curve
ECD: extracellular domain
gnomAD: Genome Aggregation Database
ICD: intracellular domain
LGIC: ligand-gated ion channels
MAFFT: Multiple Alignment using Fast Fourier Transform
P2XR: P2X receptors
PBS: phosphate-buffered saline
PISA: Protein, Interfaces, Structures and Assemblies
TEVC: two-electrode voltage clamp
TMD: transmembrane domain

## ACKNOWLEDGEMENTS

The authors thank Dr Thomas Grutter and members of the Pless laboratory for comments on the manuscript.

## AUTHOR CONTRIBUTIONS

F.G., D.S., S.B. and M.H.P. conducted functional experiments, A.S.H. performed computational analysis. F.G., D.S., M.H.P., A.S.H. and S.A.P. designed experiments and analysed data. F.G., A.S.H. and S.A.P wrote the manuscript with input from all authors. All authors have given approval to the final version of the manuscript.

## DECLARATION OF TRANSPARENCY AND SCIENITIFIC RIGOR

This Declaration acknowledges that this paper adheres to the principles for transparent reporting and scientific rigour of preclinical research as stated in the *BJP* guidelines for **<u>Design & Analysis</u>**, **<u>Immunoblotting and Immunochemistry</u>** and as recommended by funding agencies, publishers, and other organizations engaged with supporting research.

## DATA AVAILIBILITY STATEMENT

The data that support the findings of this study are available from the corresponding author upon reasonable request.

