## Supplementary information for "P2X2 receptor subunit interfaces are missense variant hotspots where mutations tend to increase apparent ATP affinity"

Running title: P2X2 receptor inter-subunit interface

Federica Gasparri[§], Debayan Sarkar[§], Sarune Bielickaite, Mette Homann Poulsen, Alexander Hauser*, Stephan Alexander Pless*

Department of Drug Design and Pharmacology, University of Copenhagen, 2100 Copenhagen, Denmark

[§] These authors contributed equally

* Co-corresponding authors:

Stephan Pless and Alexander Sebastian Hauser

Department of Drug Design and Pharmacology

University of Copenhagen

Jagtvej 160, 2100 Copenhagen, Denmark

Mail: &

Table of Contents:

Table S1……………………………………………………..………..See separate file: Table S1.xlsx

Table S2.........................................................................................See separate file: Table S2.xlsx

Figure S1……………………………………………………………………………………....02

Table S3…………………………………………………………………………………...…03

Figure S2…………………………………………………………………….........................04

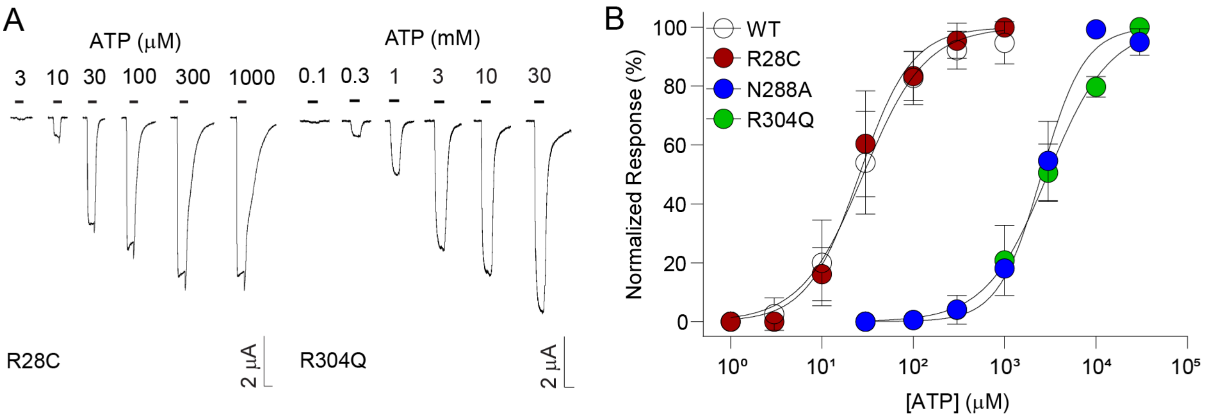


***Figure S1*: *Characterization of single point mutations present at the ATP site and from human mutation data.*** *(A) Example recordings of rP2X2R R28C and R304Q mutants. Currents are elicited by increasing concentration of ATP (black bars). Scale bar: x, 10 s y, μA. (B) Normalized ATP-elicited concentration-response data for WT and indicated mutant rP2X2Rs in response to increasing concentrations of ATP. Data are displayed as mean ± S.D. (n = 7-33).*

***Table S3:*** *List of residues lining the subunit interface of zfP2X4R that form H-bonds (HB) and/or electrostatic interactions (ION) in both conformational states (C, apo and O, ATP-bound) based on PISA analysis (default 4 Å cut-off for H-bonds and salt bridge). The atoms involved in the interactions are also shown in brackets (PDB nomenclature).* [a]*, residues known to form the ATP binding pocket therefore not analyzed further;* [b]*, interactions that were different from or not found in an earlier study* (Hausmann et al., 2014).

| Residue subunit A | Residue subunit B | Interaction type | State |
|---|---|---|---|
| S66 (OG) | R321 (NH2/1) | HB | O[b] |
| S66 (OG) | D323 (OD2) | HB | C |
| V67 (O) | R321 (NH2) | HB | O/C |
| T68 (OG1) | R321 (NH2) | HB | C[b] |
| K70 (NZ) | N296 (OD1) | HB | O[#] |
| I74 (O) | N140 (ND2) | HB | O |
| E84 (OE2) | K118 (NZ) | HB/ION | C[b] |
| R85 (NH2) | E310 (OE1) | HB/ION | O |
| I86 (N/O) | Q116 (OE1,NE2) | HB | O/C |
| I86 (O) | W167 (NE1) | HB | C [b] |
| D88 (OD1/2) | R312 (NH1/2) | HB/ION | O/C[a] |
| A90 (O) | Y302 (OH) | HB | O |
| D91 (OD1) | W167 (NE1) | HB | O |
| D91 (O/OD2) | Y302 (OH) | HB | O/C |
| D91 (OD2/1) | R312 (NH1) | HB/ION | O[a] |
| Y92 (OH) | E310 (OE1/2) | HB | C[b] |
| Q97 (NE2) | E98 (OE2) | HB | O[b] |
| D99 (OD2/1) | R321 (NH1/2) | ION | O |
| K193 (NZ) | A292 (O) | HB | O[b] |
| K193 (NZ) | G294 (O) | HB | O[b] |
| K193 (NZ) | V291 (O) | HB | C[b] |
| N195 (OD1) | N284 (N) | HB | O |
| N195 (OD1) | K285 (N) | HB | O |
| N195 (ND2) | L282 (O) | HB | C |
| R206 (NH1/2) | L282 (O) | HB | C |
| R206 (NH2) | N289 (OD1) | HB | O |
| R206 (NH2/1) | V291 (O) | HB | O |
| R206 (NH1) | A292 (O) | HB | O |
| S214 (OG) | D288 (OD1) | HB | C |
| S214 (OG) | N289 (N) | HB | C[b] |
| S214 (OG) | N290 (N/OD1) | HB | C |
| L217 (O) | R143 (NH2) | HB | O[b] |
| H219 (O) | R143 (NH1) | HB | O[b] |
| C220 (O) | R143 (NH1/2) | HB | O |
| K301 (NZ) | E310 (OE1/2) | HB/ION | O/C |
| Y303 (OH) | E310 (OE2) | HB | O/C |
| I335 (O) | Y45 (OH) | HB | C[b] |

### Western blot #1

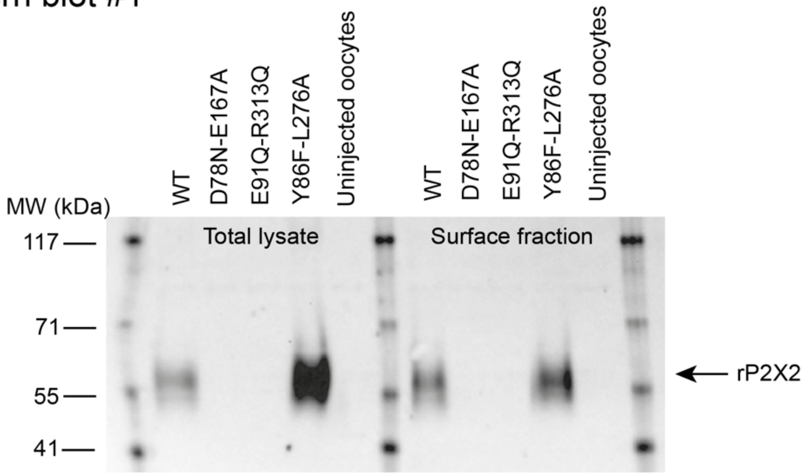


### Western blot #2

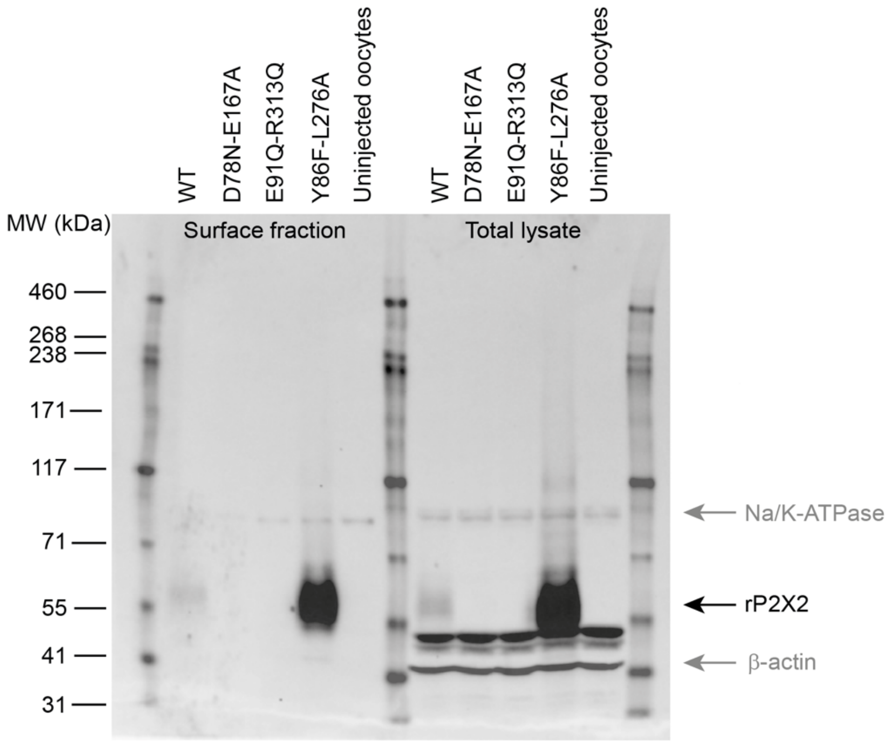


***Figure S2:*** *#1, Western blot of total lysate and surface fraction protein extracted from oocytes expressing the individual constructs (from left; P2X2R-WT, P2X2R-D78N-E167A, P2X2R-E91Q-R313Q, P2X2R-Y86F-L276A) and uninjected oocytes. #2, Western blot demonstrating that only P2X2R-WT and P2X2R-Y86F-L276A is translated into full length proteins. Na+/K+-ATPase marker was used as a loading control.*
